# Latent variable modelling and variational inference for scRNA-seq differential expression analysis

**DOI:** 10.1101/719856

**Authors:** Joana Godinho, Alexandra M. Carvalho, Susana Vinga

## Abstract

Disease profiling, treatment development, and the identification of new cell populations are some of the most relevant applications relying on differentially expressed genes (DEG) analysis. In this context, three leading technologies emerged; namely, DNA microarrays, bulk RNA sequencing (RNA-seq), and single-cell RNA sequencing (scRNA-seq), the main focus of this work. Although scRNA-seq tends to offer more accurate data, it is still limited by many confounding factors. We introduce two novel approaches to assess DEG: extended Bayesian zero-inflated negative binomial factorization (ext-ZINBayes) and single-cell differential analysis (SIENA). In addition, we benchmark the proposed methods with known DEG analysis tools for single-cell and bulk RNA data, using two real public datasets. One contains house mouse cells of two different types, while the other gathers human peripheral blood mononuclear cells divided into four types. The results show that the two procedures can be very competitive with existing methods (scVI, SCDE, MAST, and DEseq) in identifying relevant putative biomarkers. In terms of scalability and correctness, SIENA stands out from ext-ZINBayes and some of the existing methods. As single-cell datasets become increasingly larger, SIENA may emerge as a powerful tool to discover functional differences between two conditions. Both methods are publicly available (https://github.com/JoanaGodinho/SIENA, https://github.com/JoanaGodinho/ext-ZINBayes).

## 1. Introduction

Gene expression is a fundamental biological process that affects how each living organism operates. As such, studying and understanding gene expression leads to a broaden knowledge on how cells work and how they evolve. With this knowledge, ground breaking advances can be achieve in the fields of genetics, molecular biology and medicine.

One of the most relevant tasks performed through gene expression assessment is the identification of differentially expressed genes (DEG). DEG are genes that show different expression levels across different types of cells. With DEG identification we can deepen our understanding on cell differentiation, study disease phenotypes and assess how certain treatments perform [18].

Research has provided several computational methods aiming to carry out such task. Initially, differential expression (DE) analysis was only performed using gene expression obtained from DNA microarrays. Then, technological advances empowered the emergence of RNA sequencing (RNA-seq) protocols to profile gene expression. In a first approach, DE analysis over bulk RNA data was performed using packages, such as limma [21], that were initially designed to account for microarray input. However, due to differences between microarray and RNA-seq data, new methods, such as DE-seq [1] and edgeR [20], were developed specifically for the latter.

In more recent years, single-cell RNA sequencing (scRNA-seq) has stood out from the previous two. The appeal for this kind of data is the possibility to perform detailed analysis with high-resolution data, given that gene expression is described by mRNA counts in individual cells. Nonetheless, the data is still subject to the presence of noise, which unfolds as extra variation and false zero counts, caused by dropout events, batch effects, stochastic gene expression or variations in sequencing depth (or library size).

In order to prevent wrong conclusions, one must seek to disentangle correct biological information from the noisy data. One suitable approach is to use a latent variable model. Methods such as SCDE [13], MAST [7] and scVI [17] take this approach to identify DEG. However, there is a need for new techniques, since scRNA-seq datasets are becoming increasingly larger, making some of the existent methods inefficient.

In this work, we propose two new methods to perform differential expression analysis (DEA), ext-ZINBayes (extended Bayesian zero-inflated negative binomial) and SIENA (SIngle-cEll differeNtial Analysis). Both rely on a latent variable model and variational inference (VI). ext-ZINBayes adopts an existing model developed for dimensionality reduction, ZINBayes [6]. SIENA operates under a new latent variable model defined based on existing models. We benchmark their performances with other methods, using two public datasets.

In the following section, we first review latent variable models and inference concepts; then, we detail the workings of our methods (Section 2). Finally, we conclude this work with a performance analysis (Section 3) and outline some final remarks (Section 4).

## 2. Methods

As we previously mentioned, to build a scRNA method, one must account for the presence of confounding factors. Using a latent variable model has shown to be a reliable approach to separate the additional variability added by such factors.

In a latent variable model, variables are either observed or unobserved (latent). The latent variables are responsible for capturing and describing hidden factors that influence the observed variables. So, in the single cell RNA context, the observed variables would be the RNA transcript counts, and the latent would describe the confounding factors.

If we take a Bayesian perspective, i.e., if we assume that each latent variable follows a given probabilistic distribution, they can be inferred using Bayesian inference.

Under this framework, we first define probabilities that reflect a priori beliefs we may have about the latent factors. Then, these beliefs are updated using the observations which in turn generate a posteriori assumptions. This iterative process is carried out using the Bayes theorem where *p*(*Z*|*X*) reflects the a posteriori beliefs and *p*(*Z*) the a priori,

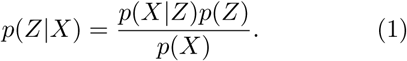

Finally, the hidden variables are inferred using the posterior, *p*(*Z*|*X*). One can set them as maximum a posteriori (MAP) estimates or as expected values of *p*(*Z*|*X*).

However, for complex models it may be impossible to obtain the exact posterior, because the marginal likelihood, *p*(*X*), can be intractable. In these cases, approximate inference techniques, such as variational inference (VI), are required.

The main idea behind VI [3] is to find a distribution *q*(*Z*) that best approximates the posterior. To do so, it assumes that *q*(*Z*) belongs to a family of distributions, defined by parameters *v*. So, in a deeper perspective, VI aims to find the parameters *v* which make *q*(*Z*) closest to *p*(*Z*|*X*). To evaluate the dissimilarity between the distributions, VI relies on the Kullback-Leibler (KL) divergence, calculated as follows:

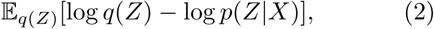

where 𝔼 _*q*(*Z*)_ is the expected value with respect to *q*(*Z*). In this setting, finding the optimal *v* amounts to finding *v* which minimize equation (2).

However, the KL divergence involves the unknown posterior, thus, an alternative metric is required. This metric is known as the Evidence Lower BOund (ELBO) and is derived from the KL divergence. The ELBO is calculated using the equation below:

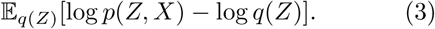

In this case, to find *v* one maximizes the ELBO.

As stated by Lopez and colleagues [17], the performance of VI techniques, is greatly influenced by the choice of the family *Q*. The most commonly used is the mean field variational family, which assumes independence between all latent variables. As such, each unobserved variable follows a separate variational distribution. Then, given a set of *N* latent variables, *q*(*Z*) can be obtained through

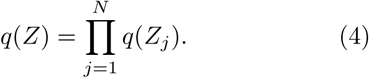

The following subsections, present two methods we developed using the techniques formerly described, with the goal to perform DE analysis. Both were developed in Python and are freely available online, ext-ZINBayes at https://github.com/JoanaGodinho/ext-ZINBayes and SIENA at https://github.com/JoanaGodinho/SIENA.

### 2.1. ext-ZINBayes

The first method we present is an extension of an existing model, ZINBayes, developed to perform dimensionality reduction. With an additional feature we enable it to detect DEG.

When designing ZINBayes, the aim was to create an approach able to discover a true biological representation of the data, without the distortion caused by noise factors. Thus, ZINBayes takes into consideration batch effects, dropout events and stochastic gene expression.

The model is build upon a Gamma-Poisson mixture, so that each count follows a Negative Binomial (NB) distribution. As research has shown, the NB is highly adequate to describe RNA-seq data due to its ability to account for overdispersion. However, it may not be sufficient to account for the excessive amount of zeros caused by dropout events. Therefore, the authors added zero inflation to the generative process.

For a given set of *G* genes and *N* cells, the count of each gene *g* in cell *i* is defined by variable *X*_*ig*_, where *g* = 1 … *G* and *i* = 1 … *N*. *X*_*ig*_ is either governed by the NB component or, in case of a dropout, is modelled as a constant zero. These conditional assignment is devised as follows,

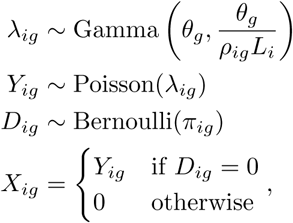

where *Y*_*ig*_ generates the count’s magnitude, if *D*_*ig*_ indicates that *X*_*ig*_ is not a dropout. *λ*_*ig*_ parameterizes *Y*_*ig*_ and thus, corresponds to the mean expression of *g* in *i*.

The latent variable *L*_*i*_ is a scale factor linked to the library size of cell *i*, i.e., the total amount of transcripts detected in cell *i*, while *θ*_*g*_ illustrates a dispersion factor associated with gene *g*. Both seen as random variables,

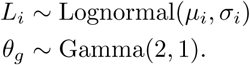

The formulations *ρ*_*ig*_ and *π*_*i*_, correspond respectively to the percentage of transcripts of gene *g* present in cell *i* and to the probability of *X*_*ig*_ being a dropout. These yield both cell-specific and gene-specific features:

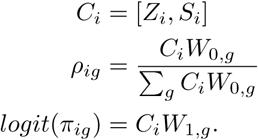

The cell-related features are the batch and the *K*-dimensional biological signature of the cell (*S*_*i*_ and *Z*_*i*_). The gene related are the factor loadings *W*_0,*g*_ and *W*_1,*g*_. While *S*_*i*_ is a *B*-sized one-hot representation, with *B* being the number of batches, *Z*_*i*_, *W*_0,*g*_ and *W*_1,*g*_ are multivariate random variables, whose components are modelled as follows,

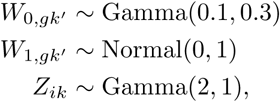

where *k* = 0, …, *K* and *k*′ = 0, …, *K* + *B*. See [6] for more details about the hyperparameters choice.

Several of the variables above are also used in scVI and have the same purpose however, scVI authors use very different parameterizations for some of them.

Given the models definition, exact inference can not be performed due to the intractability of the posteriors. In addition, the model is not conditionally conjugated, making it impossible to use coordinate ascent variational inference (CAVI). As a result, the authors resorted to reparameterization gradients (RG), a technique used in Variational Auto-Encoders (VAE) [14] and extended in Automatic Differentiation Variational Inference (ADVI) [16].

To identify DEG between two cell subpopulations we adopted the procedure developed in [17]. For each gene *g*, we define two hypotheses given a pair of cells from different populations. Yet, both cells are from the same batch and have counts *x*_1_ and *x*_2_:

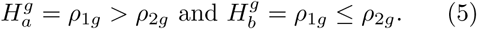

The first hypothesis states that the percentage of transcripts of gene *g* in cell 1 is higher than in cell 2, while the second hypothesis translates into the opposite. Then, a Bayes factor, *B*, is calculated as follows:

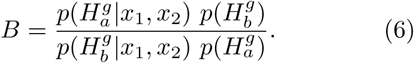

Its value quantifies the difference between the likelihood probabilities given each hypothesis. High factors reflect stronger beliefs over 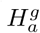, while factors closer to zero reflect more support over 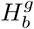. To simplify the assessment of the probability difference, we consider the factor’s logarithm and not its raw value. If the logarithm is negative it means 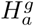 is more prone to be true, if it is positive it means the opposite: 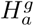 is more likely correct. This implies that higher positive values yield higher supports over 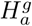, whereas lower negative values yield higher supports over the alternative hypothesis. Given that 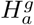 and 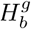 are mutually exclusive and have equal prior probabilities, i.e., 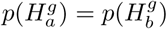, *log*(*B*) is calculated as follows:

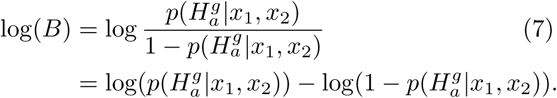

To compute the posterior 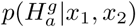, the probabilities of all *ρ*_1_ and *ρ*_2_ pairs, which make 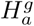 true need to be summed. Given that the *ρ* values depend on variables *Z*_1_, *Z*_2_ and *W*_0,*g*_, we need to integrate all possible combinations of those three variables that yield 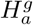 true,

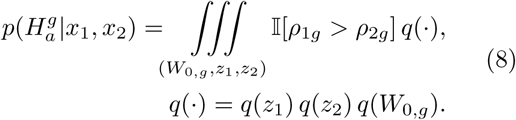

In the equation above, *q*(*z*_1_) and *q*(*z*_2_) correspond to the probabilities of cell 1 having a *z*_1_ representation and cell 2 having a *z*_2_ representation. *q*(*W*_0,*g*_) corresponds to the probability of gene *g* having *W*_0,*g*_ has its loading factors. Each of these probabilities is obtained through the corresponding variational distribution shaped during inference.

Since calculating the exact value of the integral is very computationally demanding, we used Monte-Carlo approximation. Thus, 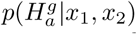 is an empirical average of *ρ*_1*g*_ *> ρ*_2*g*_ over a random set of triplets (*z*_1_, *z*_2_, *W*_0_) sampled from the variational distributions:

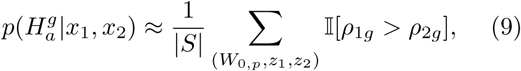

where |*S*| is the total number of samples assessed.

This process is performed over all possible cell pairs, that contain one cell from each of the two subpopulations under study. However, the *ρ* values are affected by variable *S*, which is responsible for specifying the batches and thus, this process is only viable if all cells come from the same batch. When the counts come from two or more batches, each cell must be paired with another cell from the same batch but with different type/population. If inter-batch pairs were allowed, the differences between the cell’s *ρ* could be biased by batch effects, leading to erroneous Bayes factors.

After calculating the factor’s logarithm of each pair, the obtained values are averaged and the resulting mean is used as a score of differential expression. If the absolute value of the average is higher than a certain threshold, gene *g* is classified as a DEG. The threshold used is customizable, but we recommend setting between 2 and 3 [12], since it translates into having one of the hypotheses approximately 7 to 20 times more probable than the opposite one.

To scale this procedure to very large datasets, the method enables the use of a cell pairs subset. To do so, the user needs to instruct the number of pairs to be selected. If the dataset contains cells from only one batch, we simply randomly peek the specified number of pairs. On the other hand, if the dataset gathers multiple batches, the proportion in the subset of cell pairs from each batch is equal to the proportion of each batch in the original dataset. For instance, a given dataset contains a thousand cells and 300 are from batch 1 and 700 are from batch 2. If the user specifies a subset of 100 pairs, this means that 30 of those pairs are from batch 1 whereas the other 70 are from batch 2, thus keeping the original proportions.

### 2.2. SIENA

For our second proposed method we designed a new latent variable model, where each count follows a zero-inflated NB distribution. As we mentioned before, with a ZINB distribution, one can depict the overdispersion and the excess of zero entries typical of scRNA data. Like in ZINBayes, the NB is built through a Gamma-Poisson mixture.

We decided to adopt several variables used both in ZINBayes and in scVI, making our model able to account for noise factors such as different library sizes, dropouts and stochastic gene expression. The major difference is the removal of variables *s* and *Z*, which specify the batches and the low dimensional representations of each cells biological features. Below we present the model, where *X*_*ig*_ reports the number of reads mapped to gene *g* in cell *i*:

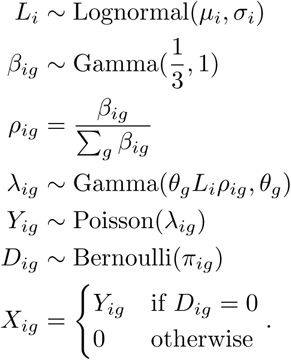

On one hand random variables *L*_*i*_, *ρ*_*ig*_, *D*_*ig*_ and *λ*_*ig*_, encode the same as in ZINBayes. *L*_*i*_ encodes a scaling factor, *ρ*_*ig*_ is the percentage of gene *g* transcripts in cell *i, λ*_*ig*_ is the expression mean and *D*_*ig*_ indicates if count *X*_*ig*_ is a dropout.

On the other hand, *π*_*ig*_, the probability of a dropout event and *θ*_*g*_, the gene’s dispersion factor are not seen as random variables; *π*_*ig*_ is a hyperparameter and *θ*_*g*_ is a non-negative model parameter. Nonetheless, these are not the only differences between this model and ZINBayes.

Similarly to what was done in [17], *L*_*i*_ is drawn from a log-normal where the mean and variance of the underlying Normal, *µ*_*i*_ and *σ*_*i*_, are set respectively as the mean and the variance of the log scaled sequencing depths/library sizes considering only cells from the same batch as cell *i*. In ZINBayes, *µ*_*i*_ and *σ*_*i*_ are the mean and variance of the log library sizes considering all cells. The choice to model *L*_*i*_ as a log-normal is to restrict its domain to be positive since its a scaling factor. Note that *L*_*i*_ encodes a factor proportionally related to the log sequencing depth, it is not the actual logarithm of the sequencing depth, as pointed out in [17].

As an alternative, we also tested *L*_*i*_ as a Gamma, where its mean and variance are equal to the mean and variance of the library sizes in *i*’s batch. In this case, *L*_*i*_ is directly related with the actual library size, and not with its logarithm.

In regards to *ρ*_*ig*_, they are set as the ratio between a factor related to gene *g* and cell *i, β*_*ig*_, and the sum of cell *i* factors with each gene. We take this formulation to not only restrict *ρ* values to be between 0 and 1, but also to constraint the sum of all *ρ*_*ig*_ of a given cell to be 1, i.e., Σ_*g*_ *ρ*_*ig*_ = 1. Both of these conditions need to be imposed because *ρ*_*ig*_ reflects a percentage, which translates into a relative frequency. An alternative approach would be to model *ρ* as a Beta distribution. However, using a Beta doesnt fit *ρ* properly since it only complies with the domain constraint. Moreover, given that no biological representation is defined for each cell, the biological variability is implicitly described directly by variable *ρ*. Notwithstanding, as it will be explained further, each *ρ* is also affected by the cells batch, since no batch-specific variable is modelled.

For the latent factors *β*_*ig*_, we chose to posit a Gamma with 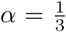 and *β* = 1 because it leads to a distribution where most of its probability density is placed near zero, yet its expected value is 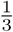. Due do its tail, this Gamma generates, in each cell, very low factors for most genes, but higher factors for a restricted set. In theory, this set is composed by cell *i* highly expressed genes.

As mentioned before, the NB is attained through a Gamma-Poisson mixture determined by variables *λ*_*ig*_ and *Y*_*ig*_, according to the following:

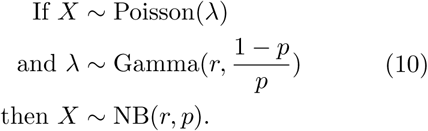

In this formulation, the NB output is defined as the number of successes until *r* failures occur, given a *p* probability of success. As a result its expected value is 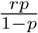. This is the NB formulation taken in our model. When deciding the parameters of the *λ*_*ig*_’s Gamma, we aimed to fix the NB expected value as *L*_*i*_*ρ*_*ig*_. By defining the Gamma’s shape as *θ*_*g*_*L*_*i*_*ρ*_*ig*_ and rate as *θ*_*g*_ we achieve that.

Finally, zero inflation is employed by variable *D*_*ig*_, which determines if *X*_*ig*_ is necessarily zero. *D*_*ig*_ is drawn from a Bernoulli distribution, since *D*_*ig*_ only needs to take two values, one indicating dropout occurrence and another one stating non occurrence. The probability of the Bernoulli, *π*_*ig*_, is set as the proportion of zero entries of gene *g* over all cells from the same type and batch as cell’s *i*. For instance, if 60% of gene *g* counts in type A and batch 1 cells are zero, then *π*_*ig*_ is set as 60% for all type A and batch 1 cells.

Regarding inference, we use reparameterization gradients. We resort to VI because the counts marginal likelihood is intractable, so exact inference can not be applied. In addition, the model is not conditionally conjugated, so CAVI can not be implemented. However, to use RG, variable *D*_*ig*_ needs to be discarded since it is not differentiable. As such, instead of defining *X*_*ig*_ with a conditional assignment, we set it as mixture of two components: one is the NB while the other models the zero-inflation, thus replacing *D*_*ig*_. The ZI part is determined by a deterministic distribution, which takes only the value zero, and the mixture proportion is set as *π*_*ig*_,

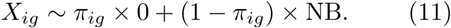

Given Equation (10), we manage to also integrate out variables *λ*_*ig*_ and *Y*_*ig*_, thus, the RG mechanism merely has to find a distribution *q* which approximates *p*(*β*_*ig*_, *L*_*i*_|*X*_*ig*_). The variational distribution *q*(*β*_*ig*_, *L*_*i*_) is considered mean-field, as such it can be factorized in *q*(*β*_*ig*_) and *q*(*L*_*i*_). Both variational distributions are assumed to be log-normal, since *β*_*ig*_ and *L*_*i*_ are positive variables and VI with reparameterization gradients performs exceedingly better when it has to optimize Normal distributions, due to the reparameterization trick.

For each log-normal we build a neural network, responsible for outputting its mean and variance, turning both *q*(*β*_*ig*_) and *q*(*L*_*i*_) into what are known as inference networks. With this approach we are able to scale inference to very large datasets, since optimization is carried only over global variables, the weights, instead of local variables, the means and variances.

Each network has one hidden layer with 128 nodes and its output layer has two heads, one for the mean and another one for the variance. A sotfplus transformation is applied over the variance head, to restrict it to be positive. In the hidden layer, a batch normalization step is employed before activation. In addition to the neural networks, memory-wise scalability is improved via batch training, where in each iteration we break the full dataset into several subsets with equal size and use each one to do an update step.

Regarding *θ*_*ig*_ optimization, we iteratively set it as a Maximum Likelihood Estimation (MLE), after one update step over the networks weights. Therefore, after optimization, *θ*_*ig*_ will have a value that maximizes the counts likelihood, given the obtained optimal variational parameters.

To assess if a given gene *g* is a DEG we apply the same procedure as the one used in ext-ZINBayes. Given a cell pair we define two exclusive hypotheses like the ones in equation (5). Then, log scaled Bayes factors are calculated for each cell pair and the absolute value of their average is used as a metric to classify *g* as a DEG or not DEG. The difference from the ext-ZINBayes procedure is the calculation of 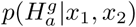, since in this approach *ρ*_*ig*_ only depends on *β*_*i*_. Consequently, it is only necessary to integrate all possible combinations of *β*_1_ and *β*_2_ that make 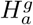 true:

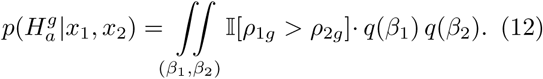

This integrals result is also approximated through Monte Carlo, where the samples are drawn from *β*_1*g*_ and *β*_2*g*_ variational distributions.

Given that in our model we do not specify any variable identifying each cells batch, the *ρ* values will be tampered by batch effects. To overcome this, we only pair cells that come from the same batch, just like in ext-ZINBayes. This way, the differential expression analysis is more truthful to biological differences. Furthermore, we scale the Bayes factor calculation by providing the optional use of a subset of pairs.

## 3. Results

To assess the performance of ZINBayes and SIENA we used two known real scRNA-seq datasets: Islam and PBMC (Peripheral Blood Mononuclear Cells), and five synthetic datasets. Since none of the real datasets has the genes identified as being DEG or not, we considered as ground truth the ones detected in the corresponding microarray dataset using limma. This is a similar procedure to the one applied in [4, 10, 13].

The Islam dataset was gathered in the study [9] and contains expression counts of 92 embryonic cells of the house mouse: 48 Embryonic stem (ES) cells and 44 Embryonic fibroblast (MEF) cells. The PBMC is a droplet-based dataset that contains count data of human peripheral blood mononuclear cells, which were sequenced in two different batches. The cells are divided in four different types, where 4996 are CD4+ T cells, 1448 are CD8+ T cells, 1621 are B cells and 339 are Dendritic cells, which amount to a total of 8404 cells.

Regarding the gold standard results, we used the microarray dataset Moliner [19] for the comparision between ES and MEF cells and two different microarray datasets of PBMC, one for the CD4+T vs. CD8+T analysis and the other for the B vs. Dendritic analysis. To obtain the Islam and the two PBMC microarray datasets (CD4+T vs. CD8+T and B vs. Dendritic) we used the GEO database [5] using the codes GSE29087, GSE8835 and GSE29618, respectively. The single cell PBMC dataset that we work with is a subset of the one used in [17]. For Moliner we extracted the data from the .CEL files^1^ used in [4].

As a preprocessing step, we filtered out the genes in the single cell datasets that were not in the corresponding microarray datasets and vice-versa. In addition, genes for which there was no information about their length were also removed, since MAST, one of the benchmarking methods, implements a TPM (Transcripts per Million) normalization, that requires the length. As such, DE analysis between types ES and MEF was carried out over 6757 genes while for the CD4+T vs. CD8+T and B vs. Dendritic analyses, only 3346 genes were evaluated.

The five synthetic datasets contain counts of 1000 genes over 1000 cells equally distributed by two conditions. Out of the 1000 genes, 200 are set as differentially expressed. To generate the datasets we resorted to the R package scDD [15], which has been already used in other studies [4] for the same purpose. More specifically, the counts were generated through scDD’s example dataset, using the *simulateSet* method. With these package we were able to devise five different gene expression scenarios according to four types of DEG, described in [4, 15]:

- traditional (DE) - unimodal gene with different expression modes in both conditions;
- DP - gene with two different expression modes shared by both conditions. However, the percentage of counts over each mode is not equal in both conditions. One has more counts closer to the first mode while the other has more counts around the second.
- DM - gene with one mode in one condition and two modes in the other, where the counts are not equally distributed. The least probable mode is equal to the unimodal condition’s mode.
- DB - combination of DP and DM types, where the cells are evenly distributed in the bimodal condition and the two modes are different from the other condition’s mode.

Using these clusters we defined four datasets corresponding to extreme scenarios. The first (200-0-0-0) has only traditional DEG, the second (0-200-0-0) has only DP differential genes while in the third (0-0-200-0) and fourth (0-0-0-200) all DEG are DM and DB, respectively. The fifth dataset contains 50 genes of each category. In all datasets, 400 of the non-differential genes are unimodal while the other 400 are bimodal. Note that in both cases the modes are the same for the two conditions. One important feature in these five datasets is that the counts are unaffected neither by dropouts (only about 10% of the entries are zeros) nor batch effects. Moreover, the differences regarding the library sizes are much smaller than in the public datasets. This means that the synthetic counts have very low noise.

In the following subsections, we first assess the effects of using different settings of SIENA, and ext-ZINBayes, then we benchmark their performances with existing methods: SCDE, MAST, scVI and DEseq. The first three were designed specifically for scRNA data whereas DEseq is used for both bulk and single-cell RNA data. To run MAST and DE-seq we used the correspoding R packages available on the Bioconductor project. For SCDE we used the R implementation^2^ provided by the authors and for scVI we used the Python release 0.3.0^3^. Finally, we compare the biological conclusions drawn from each methods DE rank through a gene set enrichment analysis (GSEA), where we compare the Gene Ontology [2] (GO) and KEGG [11] (Kyoto Encyclopedia of Genes and Genomes) pathway enrichments.

### 3.1. Configurations assessment

Inspired by what the authors in [22] concluded, we decided to evaluate how the zero-inflation affected SIENA and ext-ZINBayes. They state that to model droplet scRNA data the NB is sufficient, as such we first discuss the performance of both methods with the Islam dataset, using a simple NB or the zero-inflated version. Simultaneously, we test if the use of the gene dispersion factor improves the results. Figure 1 summarizes this analysis.

**Figure 1:**
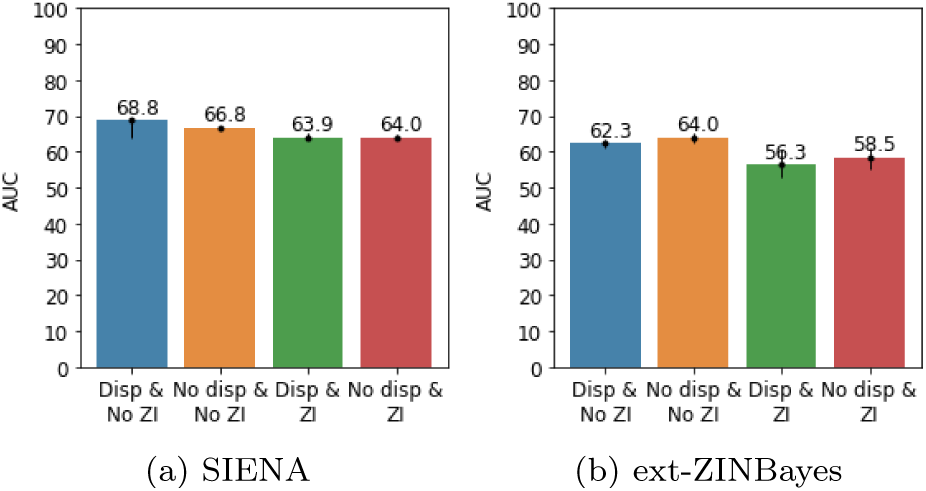
Average AUC values for each configuration with the Islam dataset. Black bars indicate the maximum and the minimum AUC achieved.

The plots show the average area under the ROC curve (AUC) for each method’s configuration: ZINB with dispersion, NB with dispersion, ZINB without dispersion and NB without dispersion. For each configuration we conducted 30 runs of 1000 epochs and averaged the resulting AUC scores. For SIENA the no zero inflation (ZI) plus gene dispersion combination yields the best average AUC, however it has the highest variance. The configurations ZI plus dispersion and ZI plus no dispersion have the lowest average. For ext-ZINBayes, employing a simple NB combined with no dispersion leads to a higher average AUC and like for SIENA, the ZI plus dispersion and ZI plus no dispersion configurations prompt the worst AUC. Yet, in ext-ZINBayes, the difference between these two configurations is more accentuated, with the former standing out has the worst. We also checked how each combination behaved with the B vs. Dendritic test from the PBMC dataset and verified what we partially concluded from Figure 1: SIENA achieves a better mean AUC with the no ZI plus dispersion combination, while ext-ZINBayes has a better mean without both ZI and dispersion. Apart from the mean variation between SIENA’s ZI configurations, all the contrasts between the plotted mean AUC are statistically significant (Welch’s t-tests with *p*-values ¡ 0.01). Therefore one can conclude that for real datasets both methods achieve better mean AUC without zero inflation, however SIENA requires the use of the gene dispersion factor whereas ext-ZINBayes does not.

For the SIENA configurations assessed, we adopted a log-normal distribution to model library scalings. However, as we mentioned in Section 2, we employed two alternatives for the library scalings, one using a log-normal distribution and another using a Gamma. From Figure 2 we see that the log-normal alternative is slightly more robust, hence our choice.

**Figure 2:**
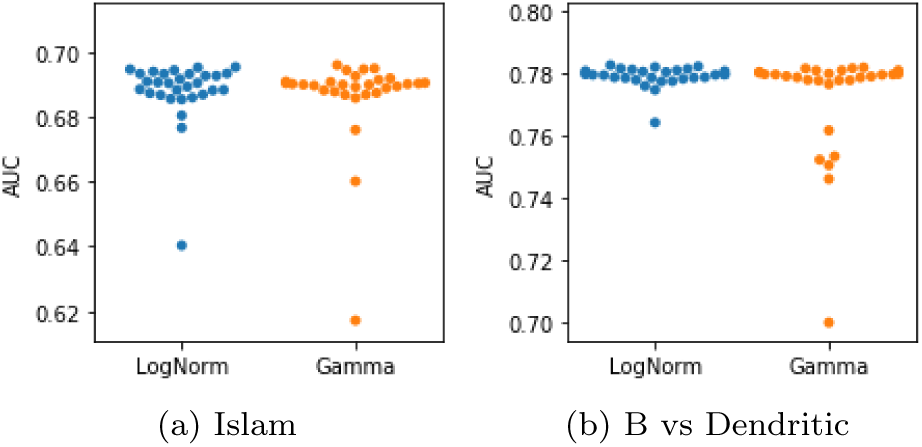
AUC values for each SIENA library size alternative. Each dot corresponds to one run.

The plots summarize the AUC values of 30 runs with each SIENA alternative, leveraging the NB plus gene dispersion configuration. Both options have similar AUC results with the Islam dataset while in the B vs Dendritic analysis, the log-normal leads to less dispersed AUC scores. In fact, the difference between the average AUC in the B vs Dendritic comparison is statistically significant (Welch’s t-test with *p*-value=0.048), however, it does not seem to be considerably high, since the resulting confidence interval at 95% is between 0.016% and 1.3%.

Given that SIENA resorts to batch training, we also used the PBMC dataset to assess how the mini-batch size affects the performance. As such, we gathered the average, minimum and maximum AUC values obtained when setting different numbers of mini-batches The results are shown in Figure 3. Although larger number of batches translate into less data used in each update step, the detection accuracy is practically unaffected by such variation.

**Figure 3:**
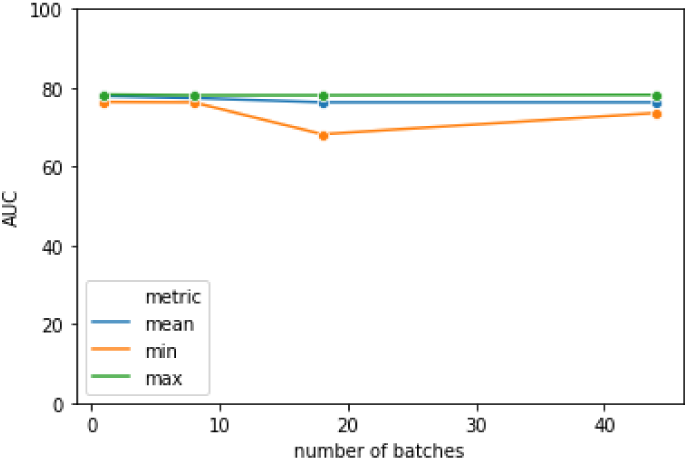
Effect of the number of batches on SIENA’s average, minimum and maximum AUC. The dots encode the metrics obtained when using 1 (no mini-batches), 8, 18 and 44, which correspond to defining mini samples of sizes 8404, 1051, 467 and 191, respectively. For each parameterization SIENA was run 30 times. Results refer to the B vs. Dendritic analysis.

### 3.2. Benchmark

To start our comparison analysis, we first contrast our methods and the four mentioned DE procedures ability to identify DEG using the Islam dataset. Out of those four only MAST and DEseq are deterministic. Similarly to what was done in the previous section, we use as a benchmark measure the average AUC. The results are shown in Figure 4.

**Figure 4:**
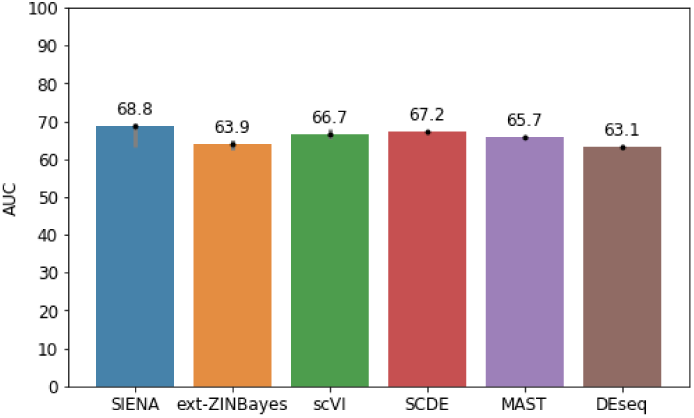
Average AUC values for each method with the Islam dataset (ES vs. MEF analysis). Grey bars indicate the maximum and the minimum AUC obtained.

To generate the bar plot, we ran and calculated the AUC of MAST and DEseq only one time, whereas for the other methods we repeated the process 50 times and averaged the AUC values. Both scVI and SIENA were run with gene dispersion and without zero-inflation, since this configuration leverages better results, and each run had 1000 epochs. ext-ZINBayes was also operated without zero-inflation and with 1000 epochs per run, but unlike for SIENA and scVI, no dispersion factor was adopted.

As seen in Figure 4, with the Islam dataset SIENA yields better results showing an average AUC close to 69%, while DEseq has the lowest average out of all the methods. SCDE has the second best AUC score, followed by scVI, MAST and ext-ZINBayes. Nonetheless, SIENA presents a higher variation (around 5%), given that two runs generated an AUC of approximately 64%. The differences between the average AUC are actually statistically significant, since all Welch’s t-tests between two methods mean AUC show *p*-values smaller than 0.01. Note that in this assessment we did not consider a t-test between MAST and DEseq since these two methods are deterministic.

With the average AUC we can test classification accuracy and robustness, however it is also pivotal to assess how each method scores the genes, i.e. how certain they are that a given gene is a DEG. As such, in Figure 5, we compare for each gene in the Islam dataset, the DE metrics of each method with the *p*-values obtained by limma. Note that the *p*-values are adjusted to the false discovery rate (FDR). For SIENA, ext-ZINBayes, scVI and SCDE we plot the metrics median of each gene considering the 50 runs conducted for the previous analysis. So, for SCDE we show the absolute *Z*-score’s median while for the other three we outline the median of the Bayes factor’s logarithm. For MAST and DEseq we only consider FDR adjusted *p*-values of one run.

**Figure 5:**
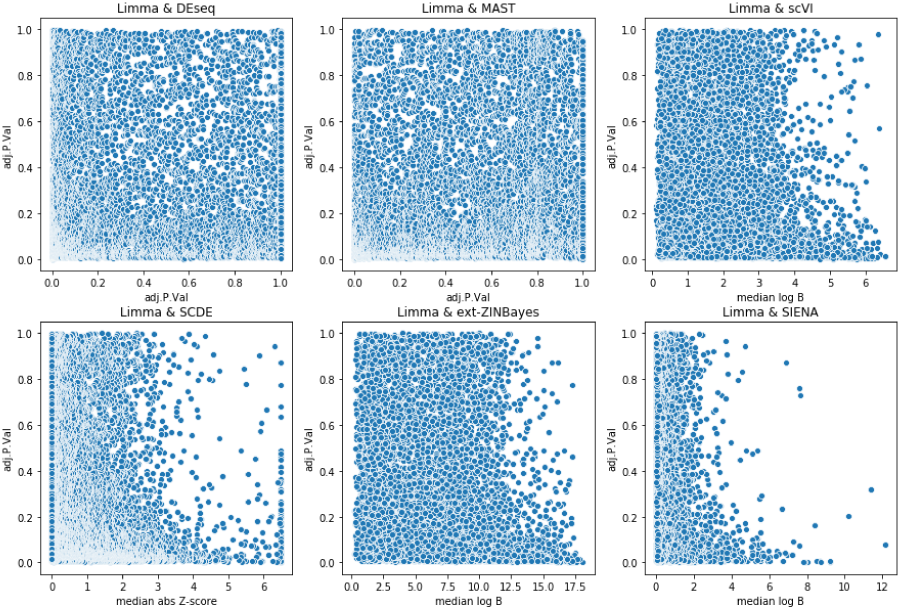
Comparison between each methods DE metrics and the adjusted *p*-values of limma. Each blue dot corresponds to a gene.

In this comparison, MAST and DEseq present the worst results since there is no visible relation between their determined *p*-values with those obtained with limma. Ideally, there should be a linear correlation. The other four methods have a more distinct correlation with limma, since at a certain threshold the *p*-values tend to be lower as the absolute *Z*-scores or log Bayes factors increase. For instance, in SCDE, genes with absolute *Z*-scores higher than ≈2.5, tend to show lower *p*-values as the score increases. Same happens for scVI, ext-ZINBayes and SIENA for genes with log Bayes factors higher than ≈ 3.5, ≈ 12.5 and ≈ 2, respectively. Nevertheless, SIENA seems to show a better correlation with limma than the others.

We also used the PBMC dataset to evaluate the methods average AUC, applying the same methodology. We also employed the same configurations for SIENA, ext-ZINBayes and scVI, but we set only 500 epochs per run. We used less iterations, because the PBMC dataset has more data entries (cells) than the Islam dataset.

As seen in Figure 6a, all methods, except SCDE and ext-ZINBayes, present a higher average AUC, when conducting DE analysis between B and Dendritic cells, than between ES and MEF cells. SCDE is the only that shows a great decrease in performance, having an average AUC lower than 50%, whereas SIENA stands out as the best with an average AUC of 77.3%. scVI and DEseq also perform well, showing a mean AUC of 76.6% and 75.9%, respectively. Unlike in the ES vs. MEF test, ext-ZINBayes shows the highest variance. Regarding the CD8 vs. CD4 comparison (Figure 6b), SIENA obtains the best mean AUC (65%), while all the other methods perform considerably worst, having an average AUC lower than 60%. Once again, SCDE shows the worst AUC. Following SIENA, MAST and scVI show, respectively, the second and third best average AUC, while ext-ZINBayes and DEseq come in fourth with an average AUC of approximately 52%. Nonetheless, similarly to the B vs. Dendritic test, ext-ZINBayes shows the highest variance in the results. Note that for this comparison, for both SIENA and ext-ZINBayes, the log Bayes factors were calculated using a subset of valid cell pairs. More specifically, 7.5 × 10^5^ pairs were used and for each of those pairs, 100 samples of *ρ* values were computed. Furthermore, some SCDE runs had to be re-executed because sometimes the method could not fit a model for a specific cell.

**Figure 6:**
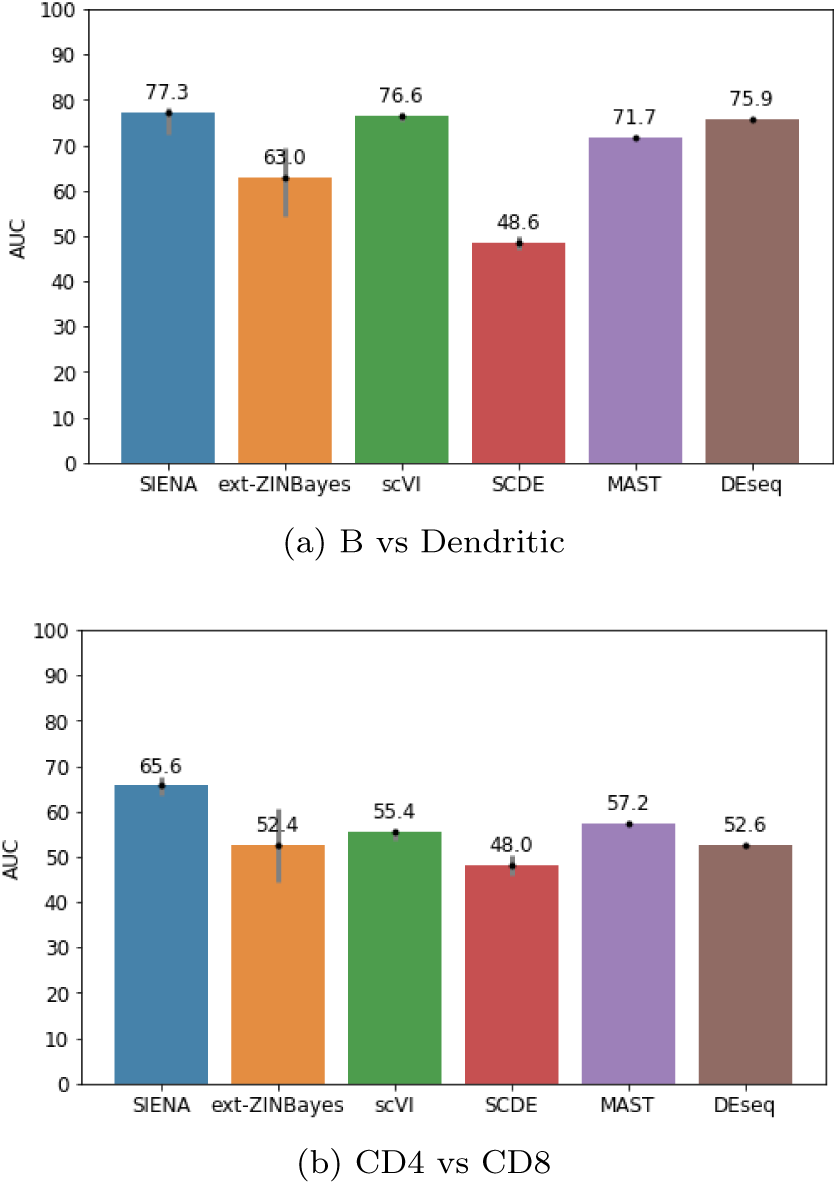
Average AUC values for each method with the PBMC dataset. Grey bars indicate the maximum and the minimum AUC achieved.

In both PBMC tests, the obtained AUC are more divergent than the ones gathered in the ES vs. MEF test. While in the latter the difference between the average AUC of the best method and the worst is slightly lower than 6%, in the B vs. Dendritic and CD4 vs. CD8 tests, the difference is around 30% and 20%, respectively.

Almost all mean AUC differences are statistically significant for both PBMC comparisons, yielding Welch’s t-tests with *p*-values lower than 0.01. The only difference that has no statistical support is between ext-ZINBayes and DEseq in the CD4 vs CD8 test (*p*-value=0.075).

Similarly to the analyses plotted in Figures 4 and 6, we collected, for each synthetic dataset, AUC results of one run for both MAST and DEseq and of 50 runs for the others methods, generating the bar charts in Figure 7. SCDE was not considered in this analysis due to the poor results achieved with PBMC and the excessive time it takes to fit the models. Unlike in the public datasets, the no zero inflation and no dispersion configuration yield better results for SIENA. As such, in Figure 7 the results regarding SIENA leverage such configuration.

**Figure 7:**
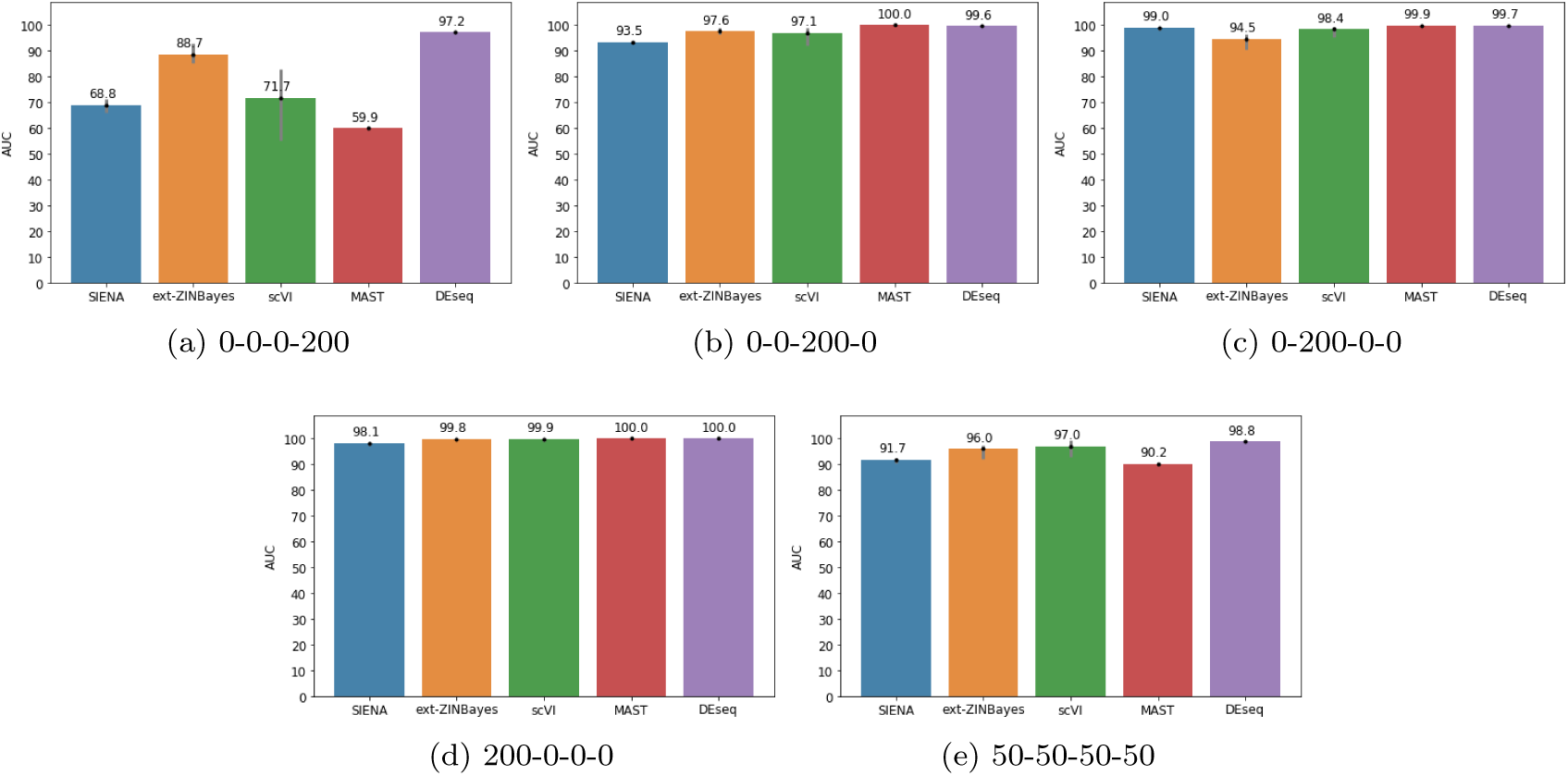
Average AUC values for SIENA, ext-ZINBayes, scVI, MAST and DEseq with each of the five synthetic datasets.

From Figure 7, we can see that in nearly all five synthetic datasets the methods yield better results, achieving average AUC higher than 90%. In fact MAST and DEseq are able to attain perfect classifications in the 0-0-200-0 and 200-0-0-0 datasets. Nonetheless, DEseq has the best performance in almost all five, leveraging AUC higher than 95%. Only in the 0-200-0-0 dataset is DEseq outperformed by another method (MAST), but only by a very small margin. Moreover, 0-200-0-0 is the sole synthetic dataset where SIENA is better than ext-ZINBayes. In all the others, SIENA is one of the two worst methods, while ext-ZINBayes is always among the top best. This contrasts with what we verified in the real datasets, where SIENA is consistently better than ext-ZINBayes.

Furthermore, with the 0-0-0-200 dataset, the performances vary substantially, prompting a 40% AUC gap between the best and the worst method. In the other four, the difference is less than 10%. Out of all methods scVI and ext-ZINBayes are the ones that show higher variations in the AUC. However, scVI stands out more due to the extensive discrepancy (around 30%) between the minimum and maximum AUC obtained with the 0-0-0-200 dataset, whereas ext-ZINBayes has a variation lower than 10% in all five.

Similarly to what we observed in the real datasets, the differences between two methods average AUC are statistically significant for all five synthetic datasets. For 50-50-50-50, 200-0-0-0 and 0-0-200-0, the differences yield welch’s t-tests with *p*-values lower than 0.05, while for the other two datasets the differences lead to *p*-values lower than 0.01.

Given the general poor results under the CD4 vs. CD8 test, we deepen our comparative analysis over the test with an intersection graph in Figure 8, without considering the microarray ground truth, i.e., the results from limma. To generate the plot, we considered the 50 runs of SIENA, SCDE, scVI and ext-ZINbayes conducted for the AUC analysis and for each method we calculated the median DE score of each gene. Then, for each method, we gathered the top 1000 genes with highest median. For MAST and DEseq, we gathered the top 1000 genes with the lowest FDR adjusted *p*-values considering only one run.

**Figure 8:**
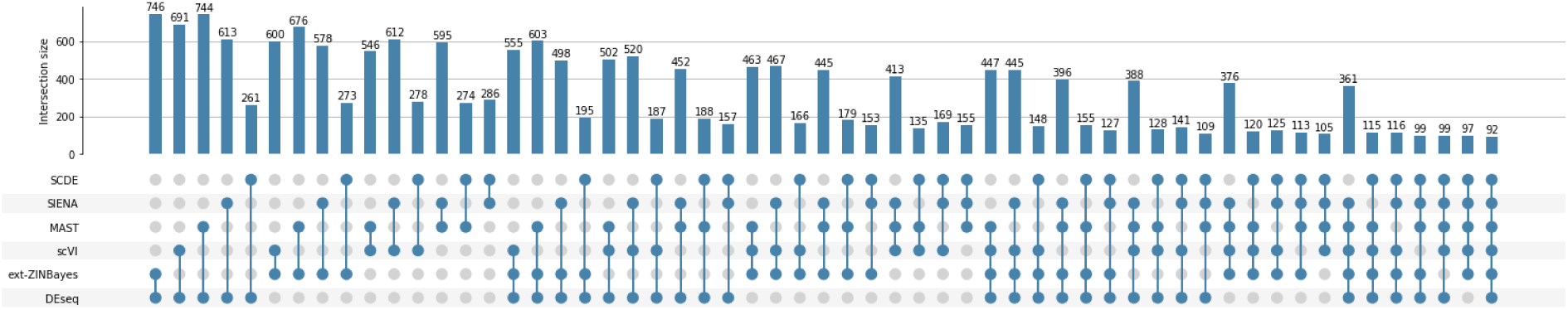
Intersections of the top 1000 DEG regarding the CD4 vs. CD8 analysis. Matrix dots specify the methods combinations and the bars encode the number of DEG in common of the corresponding combination.

From the plot, we can see that ext-ZINBayes and DEseq have a lot of genes in common, almost 750, which was expected given that the two methods have essentially the same average AUC. In fact, the pair has the largest intersection set out of all duos. DEseq has also more genes in common with SIENA than any of the other methods. This is curious given that MAST and scVI show average AUC closer to SIENA’s. Furthermore, all two method combinations considering SCDE have the lowest number of genes in common, when compared with the other two methods combinations. The same happens for three, four and five method combinations. Moreover, if we consider all methods except SCDE the number of genes in common goes from 92 to 361, it increases almost four times, whereas if one of the other methods is not considered, it only increases to values between 97 and 116. The only method that comes close to identify the same DEG as SCDE is SIENA however, the number of genes in common (287) is only a bit over 25%.

#### 3.2.1 Time

Beyond detection accuracy, it is also important to evaluate how the methods perform in terms of time usage and how that usage scales as the number of cells increases. In Table 1, we show the time that each method takes with two real data tests, MEF vs. ES and CD4 vs. CD8, and with one of the synthetic comparisons. In the first and third test the methods consider all cells in both optimization and differential expression tests, since the corresponding dataset only contains cells from the types under study. In the second, apart from SCDE, all methods take into account all PBMC entries (8404 cells) during optimization, but for the DE assessment only the subset of CD4 and CD8 cells is considered. For SCDE, the PBMC dataset could only contain CD4 and CD8 cells during the whole procedure, because it is not equipped to deal with more than two cell populations. Furthermore, in order to accurately compare SIENA, ext-ZINBayes, and scVI’s performances, only 20 Monte Carlo samples and 5 × 10^4^ pairs were considered. We take this configuration because scVI is unable to generate very large sets of *ρ* samples, due to memory over-usage.

**Table 1:** Times recorded for both proposed and benchmark methods. Results taken in a machine with a 16 core CPU and 94 GB of RAM. Synth corresponds to the 50-50-50-50 dataset.

|  | Islam | CD4vsCD8 | Synth |
| --- | --- | --- | --- |
| SIENA | 1m55s | 38m29s | 1m45s |
| ext-ZINBayes | 2m58s | 1h58m | 8m02s |
| scVI | 1m27s | 43m22s | 5m27s |
| SCDE | 5m25s | 1h29m | – |
| MAST | 1m23s | 5m54s | 27s |
| DEseq | 22s | 27s | 2m45s |

From the table, we can see that DEseq stands out as the fastest method in the real data analyses, taking less than 30 seconds. With the synthetic dataset DEseq takes longer because we had to use a different procedure to estimate a parameter (gene dispersion), than the one used for the real datasets. More specifically, a local fit had to be used instead of a parametric fit. This was necessary because the parametric fit leads to errors with the synthetic datasets. Of the two proposed methods, SIENA is the one that takes less time, yet it is still outrunned by MAST and DEseq. Nonetheless, with the synthetic dataset, SIENA is the second fastest method, taking less than half of scVI’s time. This occurs because we disabled the use of gene dispersion in the synthetic comparison. By taking this configuration, SIENA’s inference process is faster because it has one less set of parameters to optimize. In fact, if the dataset contains more cells than genes, SIENA shows better times, since it has less dispersions to optimize. This can be confirmed with the CD4 vs. CD8 comparison.

Despite fitting the model for the CD4 vs. CD8 test with only a subset of the PBMC entries, SCDE is only faster than ext-ZINBayes, taking more 50 minutes than SIENA. However, in the Islam comparison, ext-ZINBayes outperforms SCDE. This means that when SCDE considers the whole dataset, it can take more time than our two approaches.

Like scVI, SIENA and ext-ZINBayes are suitable to operate under GPUs, since both are compatible with the tensorflow-gpu library. This helps increasing their speed on machines with less processing power. Table 2 shows how SIENA and ext-ZINBayes behave in a computer with an 8 core CPU and one GPU. From the table, we can verify that the use of a GPU helps to reduce the impact of having less processing power, thus providing a suitable mechanism to make SIENA, ext-ZINBayes and scVI reach good performances in consumer-grade machines.

**Table 2:** SIENA, ext-ZINBayes and scVI Times obtained in a machine with a 8 core CPU, 1 NVIDIA GPU and 16 GB of RAM.

|  | Islam | Synth |
| --- | --- | --- |
| SIENA | 1m58s | 1m36s |
| ext-ZINBayes | 3m57s | 6m05s |
| scVI | 1m19s | 4m17s |

### 3.3. Gene set enrichment analysis (GSEA)

After gathering a DEG rank list, the next step in any differential expression analysis is to perform gene set enrichment analysis (GSEA), so as to extract biological meaning from the list. As such, it is important to compare the biological features outlined by each methods list. To do so we used the STRING [23] platform available online ^4^, to compare the gene ontologies and KEGG pathways enriched by the top DEG of each method under the B vs. Dendritic analysis. For SIENA, ext-ZINBayes, scVI, and SCDE we calculated the median DE score of each gene over the 50 runs considered for the plots in Figure 2b, similarly to what was done for the intersections plot. Then, we used the medians to rank the genes in descending order. For MAST and DEseq we used the rank list of one run.

Before feeding the lists to STRING, we first had to map the gene’s Ensembl ids to STRING ids. As a result, 5 genes were left out of the ranks because they did not map to any STRING id. Moreover, there were 6 duplicated STRING ids, thus instead of gathering 3346 items, the lists had 3347. Moreover, for MAST and DEseq the *p*-values had to be log-transformed, since STRING’s test is not sensitive enough to scores with a very large magnitude span. Due to this, 12 and 5 items were discarded respectively for DEseq and MAST, because they had a *p*-value of 0. Note that in the following analyses the SCDE method was not considered, since it had a very low average AUC with the B vs. Dendritic test.

For the ground truth (limma’s) rank, 18 GO terms were considered significantly enriched, i.e., had an enrichment FDR corrected *p*-value lower than 0.05. SIENA’s rank led to 73 enriched terms, ext-ZINBayes to 62, DEseq to 56, MAST to 41 and scVI to 31. Figure 9 illustrates for each method the enrichment score of a set of GO terms. The outlined set corresponds to the union of the 10 most significantly enriched terms by each method’s ranking.

**Figure 9:**
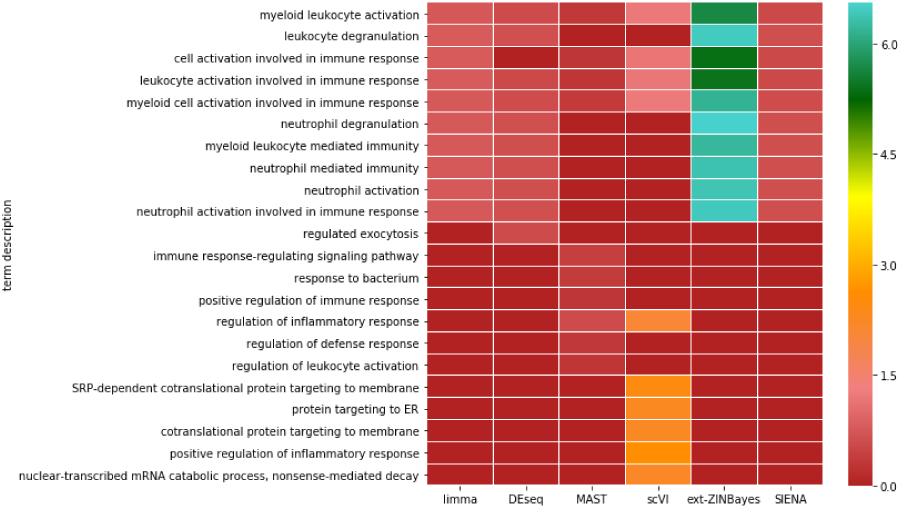
Gene Ontology enrichment analysis for each method under the B vs. Dendritic analysis. Each term considered is one of the 10 most significantly enriched terms of at least one method.

From the heatmap, we can see that the top 10 GO terms for SIENA and ext-ZINBayes ranking are the same as for the ground truth list. However, the scores related to ext-ZINBayes are greatly higher. Even though DEseq has one different enriched term, it has a closer score signature to the ground truth than ext-ZINBayes. Out of all methods, MAST shows the most divergent GO pattern. Notwithstanding, all methods seem to detect a set of DEG highly connected to biological terms such as myeloid leukocyte activation and leukocyte/myeloid cell activation involved in immune response, which means that the differences between B and Dendritic cells are probably associated with such processes.

Regarding the KEGG pathway analysis, limma led to 8 significantly enriched pathways, ext-ZINBayes to 14, DEseq also to 14, SIENA to 11, scVI to 9 and MAST to 4. Similarly to what we did for the GO analysis, we generated a heatmap (see Figure 10) showing each method’s enrichment scores for a given set of KEGG pathways. This set is the union of the top 10 significantly enriched pathways of each method.

**Figure 10:**
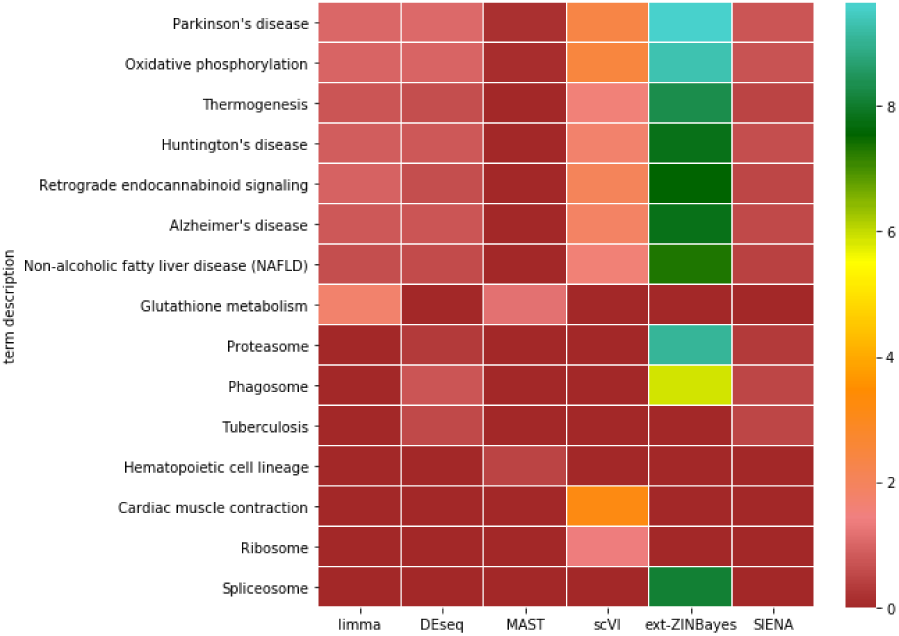
KEGG pathway enrichment analysis for each method under the B vs. Dendritic analysis. Each pathway considered is one of the 10 most significantly enriched terms of at least one method.

From the plot, we can draw similar conclusions as the ones taken from the GO analysis. SIENA, DEseq and ext-ZINBayes top pathways are very similar to limma’s top, however ext-ZINBayes enrichment scores are greatly higher. Unlike in the GO heatmap, scVI top pathways are also almost the same as limma. Moreover, all methods, with the exception of MAST, seem to be able to detect a subset of DEG related to disease pathways (non-alcoholic fatty liver and neurodegenerative diseases).

### 3.4. Discussion

In both SIENA and ext-ZINBayes, configurations without zero-inflation lead to better results in the two real datasets. This contrasts with what research has assumed throughout the years. Nonetheless, as we stated before, the authors in [22] disproved this assumption for droplet-based data, thus supporting our findings regarding the PBMC dataset. Even though the authors affirm that in the case of plate-based counts zero-inflation mechanisms are necessary, our results with the Islam counts may refute such conclusions, since that dataset has probably a plate-based origin due to its small number of cells (*<* 100).

Comparing to existing methods, SIENA was able to detect more accurately the DEG in both PBMC and Islam analysis. In addition, SIENA exhibited the most consistent behaviour over the three real data tests, showing average AUC ranging from 65% to 78%. All the other methods presented more fluctuating performances when dealing with different types of datasets. This means that SIENA is more adequate to deal with both small and large datasets than some state-of-the-art methods. Moreover, SIENA is able to scale its memory usage during both inference and DE test computation without decreasing its overall accuracy. This is important given that, in the past years, single-cell datasets have exponentially grown in size.

Contrary to SIENA, ext-ZINBayes is unable to surpass the benchmarking methods over the real data tests. In fact, in all three tests it yields the second worst mean AUC. Out of all methods assessed, ext-ZINBayes is the most unstable, reaching the highest AUC variations (around 15%) in both B vs. Dendritic and CD4 vs. CD8 tests. Given the non convexity of the loss function (ELBO), these high variations may be due to the methods inability to escape local minima when dealing with large datasets, making the methods variational parameters converge to different results in each iteration. Nonetheless with the Islam dataset, ext-ZINBayes shows a consistent behavior, whereas SIENA reaches its highest variation, which may also be due to local minima. Regarding the optimal configuration, ext-ZINBayes performs better without considering the gene dispersion factor (*θ*_*g*_). However, in both scVI and SIENA the dispersion factor improves the results, as such, we believe that a different prior or even a different formulation over *θ*_*g*_ can boost ext-ZINBayes results.

With the synthetic datasets, we observed the opposite: ext-ZINBayes identifies DEG more accurately than SIENA. In fact, it is very competitive with modern methods. Only when all the differential expressed genes are bimodal with different proportions (DP) does SIENA outperform ext-ZINBayes. This means that in most extreme scenarios ext-ZINBayes is slightly more suitable than SIENA. Nevertheless, ext-ZINBayes never yields better results than existing methods. Comparing our findings in both real and synthetic data, we can conclude that SIENA is more robust to the intrinsic noise of scRNA counts than ext-ZINBayes and other procedures. ext-ZINBayes, in turn, proves its effectiveness only with accurate data, meaning that the noise assumptions taken in its model may require adjustments.

In terms of time usage, only SIENA is able to outrun current methods in certain settings, while ext-ZINBayes is consistently one of the slowest two. This confirms what we stated before that the use of inference networks speeds up the inference process, since only global variables are optimized. Nonetheless, in the context of scRNA-seq analysis, this gain is not sufficient to make VI-based methods competitive with some alternative probabilistic procedures when it comes to time consumption.

Of the two proposed methods, SIENNA shows overall rankings more correlated with the ground truth rankings. This is not only supported by the the enrichment analysis but also by the score correlation analysis taken over the Islam dataset (Figure 5). Actually, comparing to other methods, SIENA shows DE scores more in line with limmas *p*-values. Moreover, in pair with DEseq, SIENA leads to biological conclusions closer to the ones drawn from the ground truth list.

Taking all this into account, we can conclude that only SIENA is able to compete with state-of-the-art procedures, managing to assemble more truthful differential expression scores, in a more feasible amount of time.

## 4. Conclusion

We proposed two new Bayesian probabilistic procedures to assess differential expression. Both are built upon latent variable models and variational inference mechanisms. ext-ZINBayes adopts an existing probabilistic model (ZINBayes) designed for dimensionality reduction. It performs differential analysis using some of the models latent variables. SIENA devises a novel model, leveraging certain assumptions taken in state-of-the-art methods.

Of the two procedures, SIENA yields the best results both in terms of correctness and in resource consumption. In fact, SIENA is very competitive with existing differential analysis approaches.

Both methods could benefit from time improvements. For instance, ext-ZINBayes can be upgraded with the use of inference networks whereas a new gene dispersion optimization mechanism may speed SIENA’s inference. Another potential future work, would be to integrate SIENA with some batch removal method designed specifically for scRNA data, in order to compute the Bayes factors without constraining the cells pairs by batch. One option is to employ batch correction by matching mutual nearest neighbors [8]. Finally, both models can, in principle, be used to devise some fold-change metric which can, in turn, be combined with the Bayes factor, possibly generating more accurate differential expression scores.

## Acknowledgements

This work was partially supported by national funds through Fundação para a Ciência e a Tecnologia (FCT) through projects PERSEIDS (PTDC/EMS-SIS/0642/2014), PREDICT (PTDC/CCI-CIF/29877/2017) and NEUROCLINOMICS2 (PTDC/EEI-SII/1937/2014), and under contracts IT (UID/EEA/50008/2013), INESC-ID (UID/CEC/50021/2019) and LAETA (UID/EMS/50022/2019).

## Footnotes

1 http://carlosibanezlab.se//Data/Moliner_CELfiles.zip

2 https://hms-dbmi.github.io/scde/package.html

3 https://github.com/YosefLab/scVI/releases

4 https://string-db.org

